# Detecting and quantifying changing selection intensities from time-sampled polymorphism data

**DOI:** 10.1101/027961

**Authors:** Hyunjin Shim, Stefan Laurent, Matthieu Foll, Jeffrey D. Jensen

**Affiliations:** School of Life Sciences, Ecole Polytechnique Fédérale de Lausanne (EPFL), Lausanne, Switzerland; Swiss Institute of Bioinformatics (SIB), Lausanne, Switzerland; International Agency for Research on Cancer, Lyon, France

**Keywords:** population genetics, fluctuating selection, change point analysis, time-sampled data, approximate Bayesian computation, Wright-Fisher model, experimental design

## Abstract

During his well-known debate with Fisher regarding the phenotypic dataset of *Panaxia dominula*, Wright (1948) suggested fluctuating selection as a potential explanation for the observed change in frequency. This model has since been invoked in a number of analyses, with the focus of discussion centering mainly on random or oscillatory fluctuations of selection intensities. Here, we present a novel method to consider non-random changes in selection intensities using Wright-Fisher approximate Bayesian (ABC)-based approaches, in order to detect and evaluate a change in selection strength from time-sampled data. This novel method jointly estimates the position of a change point as well as the strength of both corresponding selection coefficients (and dominance for diploid cases) from the allele trajectory. The simulation studies of CP-WFABC reveal the combinations of parameter ranges and input values that optimize performance, thus indicating optimal experimental design strategies. We apply this approach to both the historical dataset of *Panaxia dominula* in order to shed light on this historical debate, as well as to whole-genome time-serial data from influenza virus in order to identify sites with changing selection intensities in response to drug treatment

## Introduction

The common assumption of constant selection intensity through time utilized in many tests of selection is often criticized as unrealistic in natural and experimental populations - both owing to environmental changes (e.g., fluctuations in climate, predation, or nutrition) as well as to genetic changes (e.g., epistasis, clonal interference). Despite this, such considerations are not accounted for in most population genetic models, since inferring changing selection coefficients (*s*) from single-time point polymorphism data is difficult. However, owing to recent technological advances, time-sampled polymorphism data are increasingly available, and time-serial analytical methods are expanding (Malaspinas et al. 2012; Mathieson and McVean 2013; Foll et al. 2014a; Lacerda & Seoighe 2014; and see review of Bank et al. 2014) – allowing for an empirical evaluation of the importance of changing *s* models.

Fluctuating selection in natural populations was suggested by Wright (1948) with regards to the phenotypic time-serial data of *Panaxia dominula* (scarlet tiger moth) to account for its observed annual fluctuations (Fisher and Ford 1947). Since, there have been several theoretical considerations of fluctuating selection (Kimura 1954; Karlin & Levikson 1974; Karlin & Lieberman 1974; Gossmann et al. 2014; Gompert 2015), as well as many observations of fluctuating selection in natural populations (for a review, see Bell 2010). Nonetheless, until recently, analyses of fluctuating selection centered on *random* or *seasonal* oscillations of selection strength through time, as the mathematical complexity of analytical methods only allowed the simplest cases to be considered.

Approximate Bayesian Computation (ABC) has the advantage of being flexible in integrating complex models due to computational efficiency and the lack of likelihood computation (Beaumont 2010). Recently, a hierarchical ABC-based method based on the Wright-Fisher model was developed in order to infer genome-wide effective population size and per-site selection coefficients from whole-genome multiple-time point datasets (Foll et al. 2014a, b). While the initial approach performs well overall, the authors noted the possibility for observations inconsistent with a single-*s* Wright-Fisher model; this was indeed observed at certain sites in their analysis of the influenza virus genome. In their analyses, these trajectories are simply excluded from consideration. Thus, as a natural extension, we here investigate the presence of changing selection in these outlier SNPs; in doing so, we also develop an extended Wright-Fisher ABC-based method capable of detecting and quantifying changing selection intensities through time.

## Materials and Methods

### Wright-Fisher ABC-based method

This approach first relies on a previously developed Wright-Fisher ABC method (WFABC; Foll et al. 2014a,b) in order to estimate effective population size (*N_e_*). The posterior of the *N_e_* estimated from WFABC is used as a prior for the following extended method. The trajectory *X* of a given allele with a known *N_e_* consists of time-serial allele frequencies f*_t_* (*t* = 1,…,*T*) where *T* is the total number of generations (with *T* > 4 to allow for a change point to be realizable with the Wright-Fisher model), from which a sample *n_i_* is taken at sampling time points *i =* 1,…,*I* (with *I* > 4 to allow for a change point to be detectable). Parameters to be inferred include the selection coefficient prior to the change in selection intensity (*s_1_*), the selection coefficient subsequent to the change in selection intensity (*s_2_*), the time of change (*CP*), and the dominance coefficient (*h*) for diploid models. The joint posterior distribution of these parameters can be estimated by

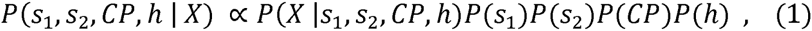

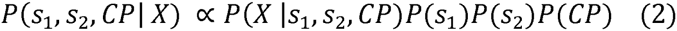

for the diploid model and the haploid model, respectively. The ABC approach allows these parameters to be inferred using Wright-Fisher model simulations without calculating the likelihood *P*(*X*|*s*_1_*, s*_2_, *CP, h*) or *P(X*|*s*_1_, *s*_2_, *CP*).

The Wright-Fisher model simulator with a change point in selection strength is used to simulate the data *X,* with relative fitnesses *W_AA_=*1+*s, W_Aa_*=1*+sh* and *w_aa_*=1 for the diploid model, and *W_A_=1+s* and *w_a_=1* for the haploid model (Ewens 2004). Initially, the random sampling of an allele from generation *1* to generation *CP-1* is simulated using *s_1_*, and onwards from the change point (*CP*) using *s_2_*. In order to simulate realistic allele trajectories with changing selection coefficients, the allele needs to be segregating at the time of the change point. This condition is necessary since the change in selection coefficient cannot occur if the allele is either lost or fixed beforehand, assuming the infinite-site model with no back mutations. Thus, only alleles segregating at the change point are accepted as a data censoring procedure.

The associated summary statistic for these time-serial data is *Fs’,* an unbiased estimator of *N_e_* that measures the allele frequency change between two sampling time points without bias in cases of highly skewed allele frequencies and cases of small sample size (Jorde and Ryman 2007). It is given as

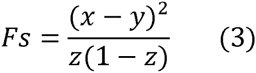

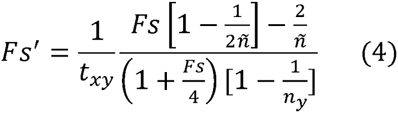

where *x* and *y* are the allele frequencies at two consecutive time points separated by *t_xy_* generations, *z* = *(x+y)/2,* and *ñ* is the harmonic mean of the chromosome sample sizes *n_x_* and *n_y_* at two consecutive time points. Unlike the WFABC approach that summarizes time-serial trajectories into only two summary statistics (increasing and decreasing *Fs*’; Foll et al. 2014a), here *Fs’* is summarized at every pair of consecutive time points as *Fs’_1_,…,Fs’_1-1_,* where *I* is the number of sampling time points. This modification allows additional information such as the timing of increase or decrease in allele frequency to be captured - an important factor for detecting the change point. In order to retain information about directionality, increasing allele frequencies are made positive and decreasing allele frequencies are made negative with regards to the absolute value.

The joint posterior distribution of the parameters of interest is obtained using the algorithm described in Beaumont et al. (2002). The approximate posterior density

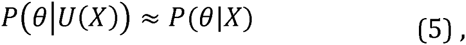

with θ=(*s_1_*, *s_2_, CP, h*) for the diploid model and θ=(*s_1_*, *s_2_, CP*) for the haploid model, is obtained using an ABC algorithm as follows:

i. Simulate *K* trajectories from the Wright-Fisher model with a change in selection intensity, with *θ* randomly sampled from its prior P(*θ*), conditional on the allele segregating at the change point.
ii. Compute U(*x_k_*) for each accepted trajectory using the *Fs*’ summary statistic between all consecutive sampling time points *i*: U*(x_k,i_)* where *i* = 1*,…,I-1* where *I* is the last sampling time point.
iii. Retain the simulations with the smallest Euclidian distance between U*(x_k,i_)* (from the simulated) and U*(X_i_)* (from the observed) to obtain an approximate posterior density of P*(θ/X).*

For the first step, simulations are performed with the same initial conditions as the observed data - including effective population size, initial allele frequency, and the sampling points and sizes. In addition, a minimum allele frequency in one of the sampling time points is imposed on simulated trajectories as is done in observed data. This ascertainment scheme takes into account the non-random criterion of considering only the trajectories reaching values above the sequencing error threshold in the observed data (Foll et al. 2014b).

For the second step, it is important to note that the *Fs’* summary statistic is calculated between every pair of consecutive sampling time points (Figure 1) – thus there are *I*-1 summary statistics for each simulated and observed trajectory. This construction of the summary statistic enables information on both the timing and strength of the allele frequency change to be captured, as the timing of the change is essential in detecting the change point and the strength of the change is essential in estimating the corresponding selection coefficients. For the diploid model, an additional parameter *h* is inferred jointly with the other three parameters, as its value is one of the determining factors in the timing of allele frequency change (Haldane 1932).

**Figure 1.**
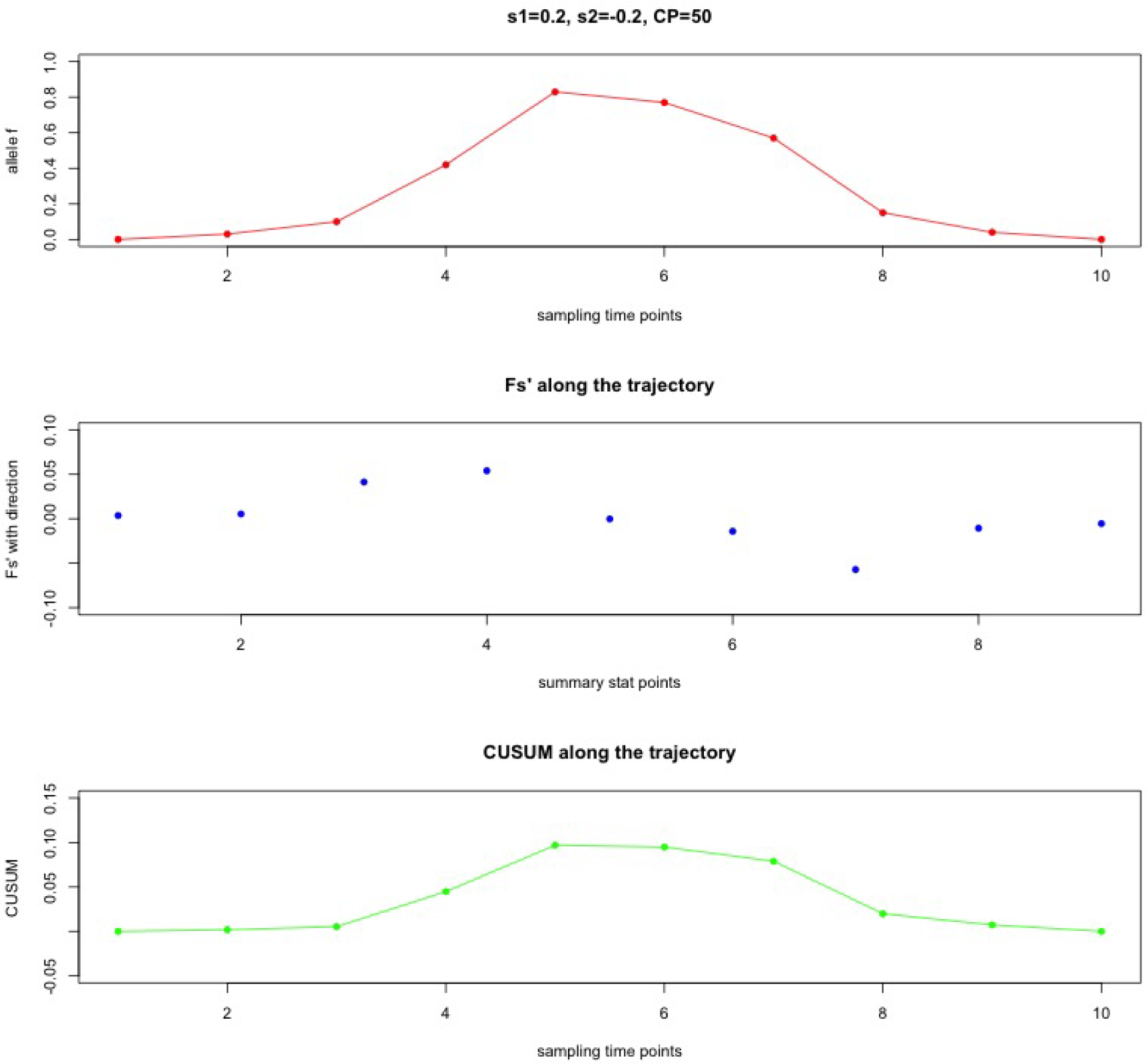
Illustration of *Fs’* calculated between every pair of consecutive sampling time points and the maximal CUSUM value *S_cp_* as summary statistics, using a haploid population of *N_e_ =* 1000 with a *de novo* mutation and the sample size as 100.

For the third step, the simulated *Fs*’ summary statistics U*(x_k,i_)* between every pair of consecutive sampling time points are compared with the corresponding observed *Fs’* summary statistics U*(X_i_) –* allowing a small fraction of the simulated trajectories (less than 0.1%) with allele frequency changes that best match the observed trajectory (in terms of both timing and strength) to be retained.

### Wright-Fisher ABC-based method with Change-point analysis

In order to increase computational efficiency and sensitivity in change point detection, an additional summary statistic is integrated into the Wright-Fisher ABC-based method. This novel summary statistic is derived from change point analysis – statistical techniques developed and used in many disciplines ranging from finance to quality control in order to detect and estimate change (e.g., Chen and Gupta 2001). Among the techniques available, the cumulative sum control chart (CUSUM) developed by Page (1954) is able to detect small and sustained shifts in the statistics *β* obtained from a sample (Ryan 2011). Instead of using the entire CUSUM procedure as a separate method for detecting change, the CUSUM value is integrated into the Wright-Fisher ABC-based method as an additional summary statistic that characterizes the time-sampled trajectory of an allele:

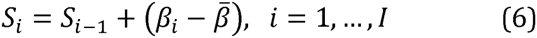

where 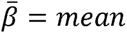 and *S*_0_ = 0. The CUSUM value *S* is accumulated only when the statistic *β* is different from its average value in the dataset.

The change point *S_cp_* is the sampling time point with the maximal absolute value of *S_m_,* which is the furthest point from the initial value zero attaining the maximal accumulation of difference from the average value:

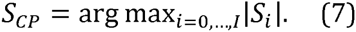

Here, we calculate *Fs’* at each pair of consecutive sampling time points as the statistic *β,* since it is a time-serial measure of the allele frequency change - which is indicative of the selection strength change. Thus, when *Fs’* is used as the statistic *β* in the CUSUM, the maximal CUSUM value *S_cp_* is the potential change point of the allele trajectory, as illustrated with an example in Figure 1.

In the Change-Point Wright-Fisher ABC (CP-WFABC), an additional summary statistic *S_cp_* with an infinite weight is used to characterize observed and simulated allele frequency trajectories for detecting a change point In the third step of the ABC algorithm, the Euclidean distance between U(*x_k,i_)* and U*(X_i_)* is calculated only if the maximal CUSUM value *S_CP,k_* of the simulated data matches the maximal CUSUM value *S_cp_* of the observed data:

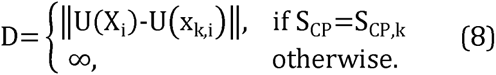

This additional step allows the computation to be more efficient – especially when there is a large number of time points sampled – as the Euclidean distance is calculated for a fraction of simulated trajectories whose maximal CUSUM value is equal to that of the observed (i.e., with the same time-sampled characteristic). Furthermore, as the CUSUM is sensitive to small and sustained changes, integrating the CUSUM into the Wright-Fisher ABC increases its sensitivity for detecting small and sustained changes in selection strength. The potential bias in the calculation of the maximal CUSUM value is counteracted by the fact that the bias would be present in both the observed and the simulated trajectories.

### Simulated data with constant selection and with changing selection

We generated simulated datasets of different effective population sizes using the Wright-Fisher model for two scenarios: (1) trajectories of constant selection with only *s* and *h* (for diploid models) as parameters, and (2) trajectories of changing selection with *s1, s2, CP,* and *h* (for diploid models) as parameters. For selection coefficients, uniform priors of [-1,1] were used. The uniform prior of *CP* was set to occur between the second generation and the second-to-last generation [2,*T*-1], where *T* is the number of generations of the population in the time-serial data. The dominance coefficient *h* for the diploid model was randomly drawn from one of three values: complete recessiveness, co-dominance, or complete dominance [0,0.5,1]. Although these prior ranges are uninformative, the constraint on the trajectories to be segregating at the change point shapes the distribution of the prior ranges according to the input parameters such as ploidy, effective population size, initial allele frequency, and number of generations; the updated priors for the haploid population of *N_e_* = 100 and the diploid population of *N_e_* =50 are shown as examples (Figure S1 and S2).

The other input values – such as the number of generations (*T*=100), the sampling time points (*I*=10), the sample size (*n*=100), the initial allele arising as a new mutation, and the ascertainment of observing a minimum frequency at 2% – were kept constant for the two scenarios. We retained the best 0.1% of 1,000,000 simulations for each pseudo-observable trajectory using the rejection algorithm based on the Euclidean distance as described above. The mode of the posterior distribution from the best simulations (Sunnåker et al. 2013) was used to evaluate the estimated parameter value against the true parameter value.

## Results

### ABC model choice in the Change-Point Wright-Fisher ABC method

The first step of CP-WFABC is to be able to distinguish changing selection trajectories from constant selection trajectories. ABC model choice was constructed to choose between two models: M_0_ with a single selection coefficient, and M_1_ with two selection coefficients and a change point The relative probability of M_1_ over M_0_ can be computed through the model posterior ratio as the Bayes factor *B_1,0_* (Sunnåker et al. 2013):

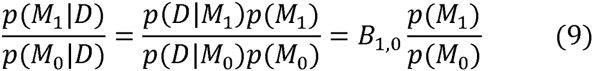

when the model prior *p*(M_0_) is equal to *p*(M_1_). In practice the model priors are made equal by producing the same number of simulations for each model and retaining the best simulations from the lot. The posterior ratio is computed as the number of accepted simulations from M_1_ over those of M_0_ – giving the Bayes factor *B_1,0_* which is an indicator of the support for a specific model. The performance study was conducted with a haploid population of *N_e_ =* (100,1000, or 10000) and a diploid population with *N_e_ =* (50, 500, or 5000) using the simulated datasets of the two scenarios described in the previous section as M_0_ and M_1_, respectively.

We considered two cases for the pseudo-observables to test the sensitivity and specificity of the ABC model choice: the first case when the pseudo-observed trajectories have a single selection coefficient, and the second case when they have changing selection coefficients with a change point. One thousand pseudo-observable trajectories were generated for each case with the data ascertainment minimum frequency set to 2% for at least one of the sampling time points. Additionally for the second case, pseudo-observable trajectories were accepted only when the allele was segregating at the time of the change point – a constraint for realistic combinations of selection coefficients, change points, and dominance (for diploids) – in order to reproduce changing selection trajectories in real datasets. All other input values were kept constant as in the simulated datasets described in the previous section.

The results of the ABC model choice from a haploid population with *N_e_ =* 100 and a diploid population with *N_e_ =* 500 are represented as ROC curves (Robin et al. 2011) in Figure 2. Specificity is given on the x-axis showing the true negative rate, while sensitivity is given on the y-axis showing the true positive rate of the Bayes factor *B_1,0_* calculated from 1000 pseudo-observables of changing selection (where *B_1,0_* should be large) and 1000 pseudo-observables of constant selection (where *B_1,0_* should be small). The overall ROC curves in black (all trajectories) show that when the specificity threshold is most conservative in detecting no false positives (i.e. *B_1,0_ =* infinite), the Bayes factor *B_1,0_* has a sensitivity of around 30% for all populations. Considering that the pseudo-observable trajectories were simulated randomly from a wide range of prior values, the Bayes factor *B_1,0_* from CP-WFABC is in general sensitive and specific. The ROC curves in black for the other haploid and diploid populations (Figure S3) also indicate that the Bayes factor *B_1,0_* is sensitive and specific as they are above the diagonal line of no-discrimination. The area under the ROC curve (AUC) is used to assess how reflective the Bayes factor *B_1,0_* is of the true model, as summarized for all pseudo-observable populations in Table 3. The AUC values show that the Bayes factor *B_1,0_* is ~80% more probable to rank a randomly chosen changing selection case above a randomly chosen constant selection case. Additionally, the distribution of Bayes factors *B_1,0_* under the null model M_0_ (i.e., case 1) was used to compute the significance level α at 1% (Good 1992). For both diploids and haploids, the significance threshold is higher for smaller population sizes (Table 4), and the calculation of these thresholds will be important in any given data application.

**Figure 2.**
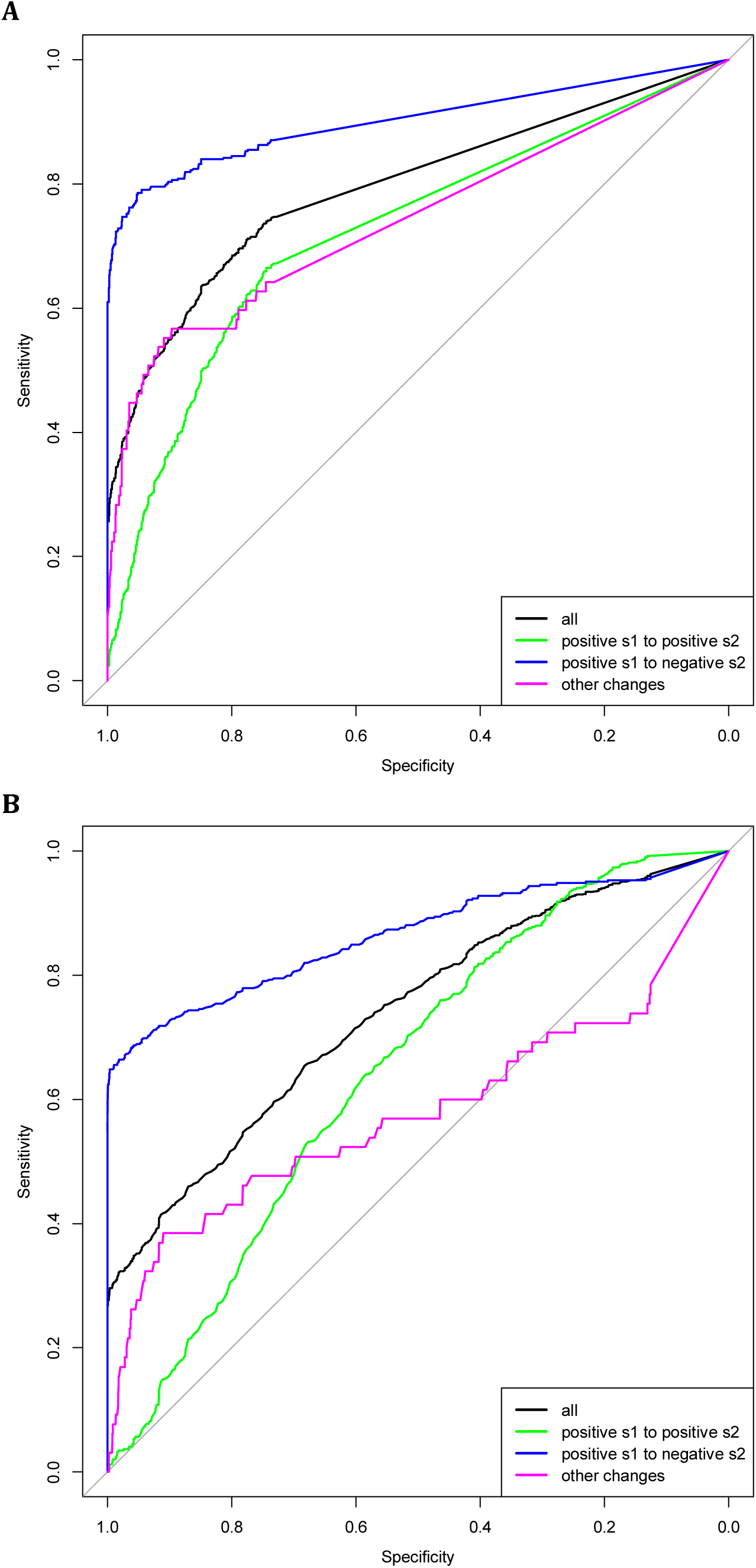
ROC curve of the Bayes factor *B_1,0_* from the ABC model choice of a haploid population with *N_e_*=100 (A) and a diploid population with *N_e_=* 500 (B).

Following the detection of changing selection trajectories using ABC model choice, the quality of parameter estimation by the model chosen was evaluated. The cross-validation results from the haploid population of *N_e_ =* 100 are shown in Figure 3 and those from the diploid population of *N_e_* = 500 in Figure 4 (see Figures S4-S7 in Supporting information for additional results). For the case where the pseudo-observables were of constant selection, the estimation for a single *s* (and the dominance *h* for diploid) using CP-WFABC is very accurate, as the mode of the best simulations from the M_0_ model for each pseudo-observable lies along the red diagonal line. Exceptions include uninformative trajectories where the allele surpasses the minimum frequency of 2% in the first sampling and is lost immediately due to genetic drift or negative selection and therefore not observed in subsequent samplings. Such trajectories will always keep the same set of best simulations from the M_0_ model since their selection strength is indistinguishable, and they result in horizontal lines along the estimated negative value. This phenomenon is particularly pronounced when population size is small as shown in Figures 3 and S6, since the role of genetic drift is more significant.

**Figure 3.**
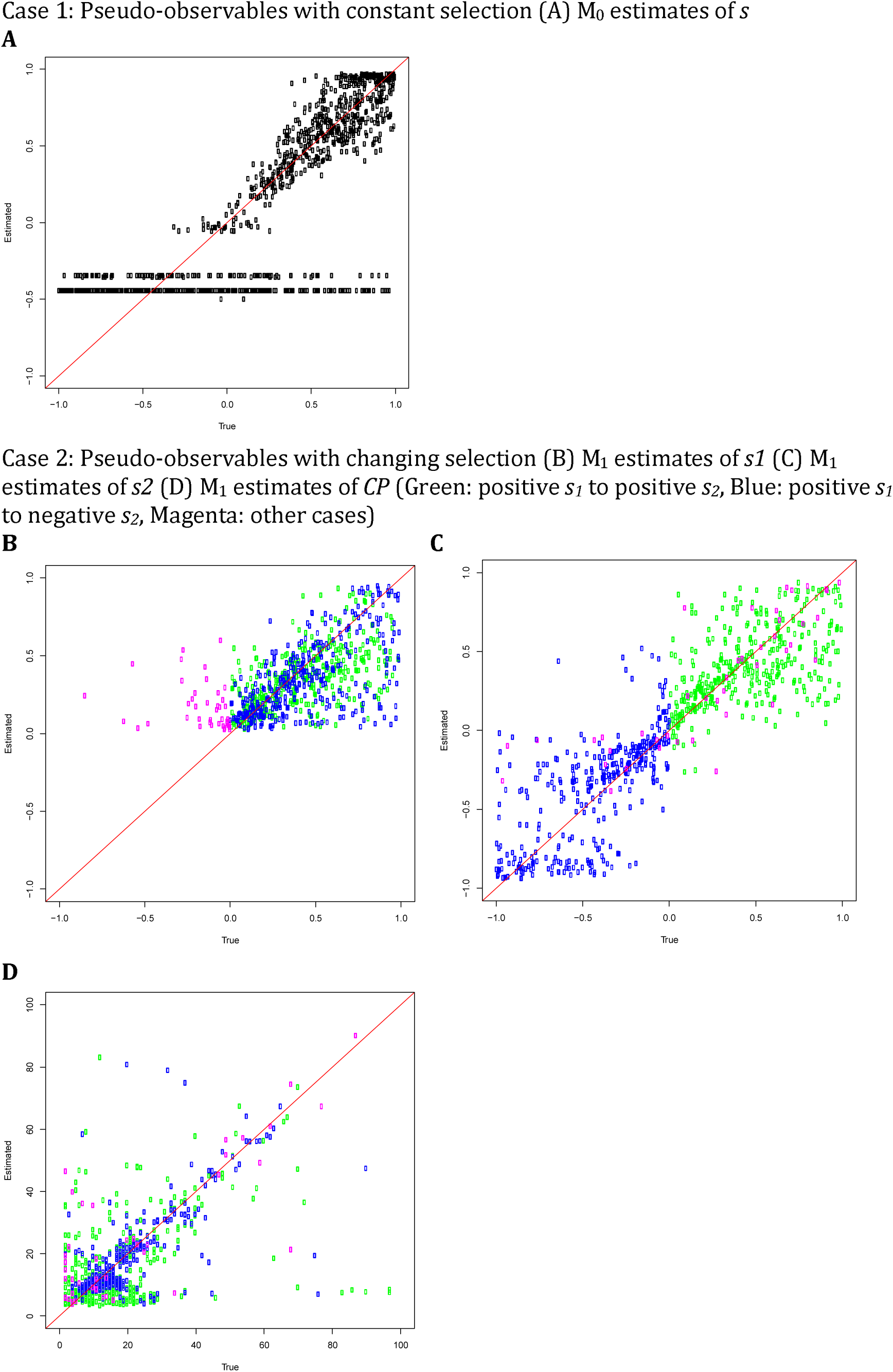
ABC model choice parameter estimations for **1000** pseudo-observables with a haploid population of *N_e_*=100. Each circle is the mode of the posterior distribution from the 0.1% best simulations.

**Figure 4.**
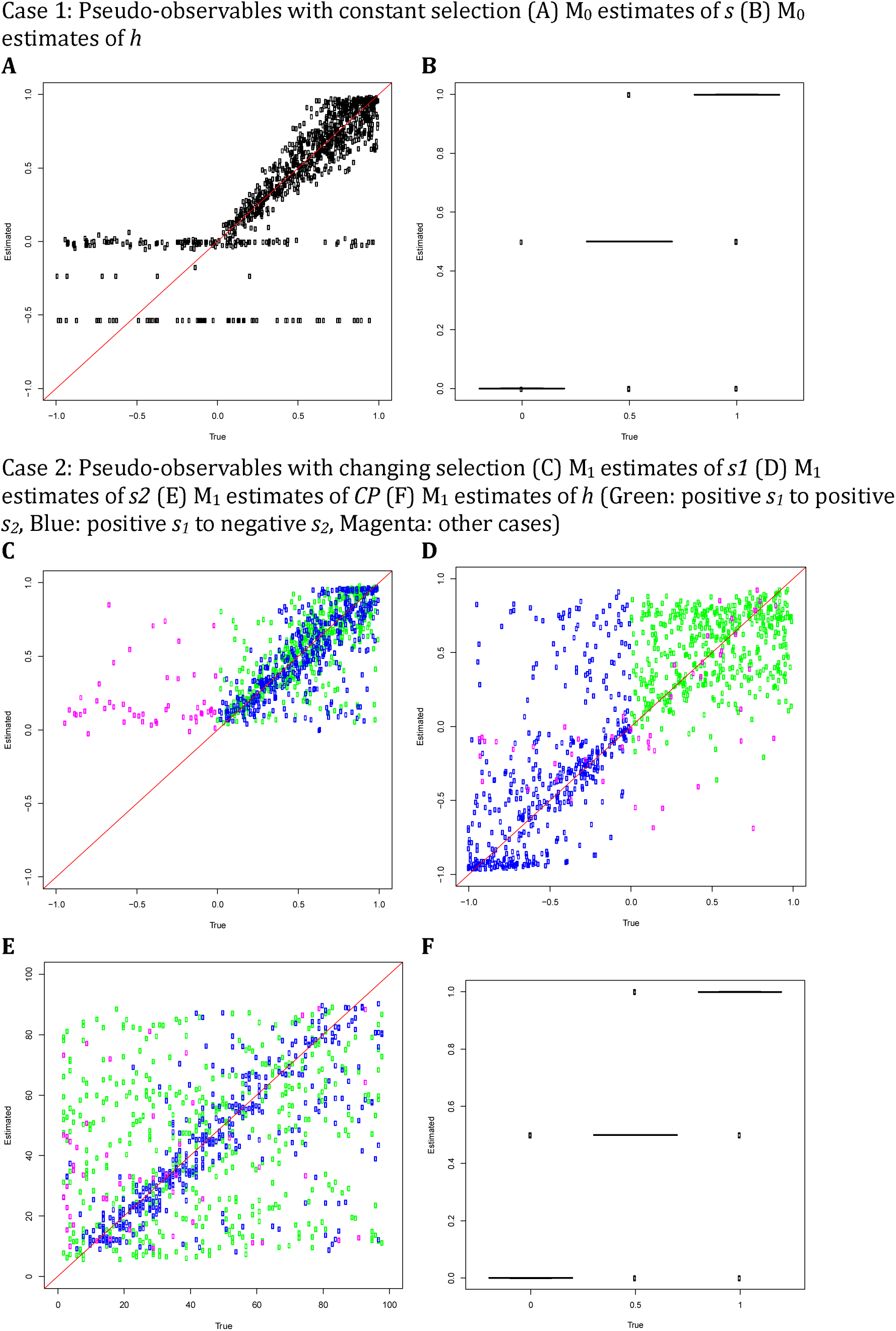
ABC model choice parameter estimations for 1000 pseudo-observables with a diploid population of *N_e_=* 500. For cross-validation graphs, each circle is the mode of the posterior distribution from the 0.1% best simulations. For boxplots, red dots are true values and blue dots are average estimated values.

For the second case when the pseudo-observables are of changing selection intensity, the joint estimation of the parameters is also effective for a restricted range of values. In Figures 3-4 and S4-S7, the mode estimation of each pseudo-observable is color-coded according to the three categories of trajectory shape. The green dots are pseudo-observable trajectories that change from positive *s_1_* to positive *s_2_*. The blue dots are those that change from positive *s_1_* to negative *s_2_,* while the magenta colors include all other cases (e.g., neutral or negative *s_1_* to any value of *s_2_*). There is a clear clustering by category – with the best estimation being of positive values of *s_1_* below 0.5, moderate values of *s_2_* between -0.5 and 0.5, and CP values for the blue category of positive *s_1_* to negative *s_2_.* In trajectories other than those with positive *s_1_* to negative *s_2_,* the change point is difficult to detect, particularly for diploid populations where the additional dominance parameter *h* was estimated (Figure 4 and Figure S6-S7). This trend is also observed when the ROC curves are generated according to these three categories (Figure 2 and Figure S3). For all populations, the Bayes factor *B_1,0_* is more sensitive and specific for trajectories changing from positive *s_1_* to negative *s_2_* (ROC curves in blue), reaching above 60% of the true positive rate when there are no false positives. Despite the restricted range of good parameter estimation in *s_1_, s_2_* and *CP,* the estimation of dominance is robust for both cases of constant and changing selection (except for the small population size of *N_e_* = 50; Figure S6).

In order to evaluate the performance of the joint parameter estimation, the coefficient of determination *R^2^* is used to assess the cross-validation between the estimated values (*y_m_*) and the true values (*f_m_*), compared with the simple average of the estimated values 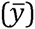. The closer the *R^2^* value is to 1, the better the parameter estimation as shown:

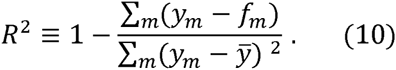

**Table 1.**
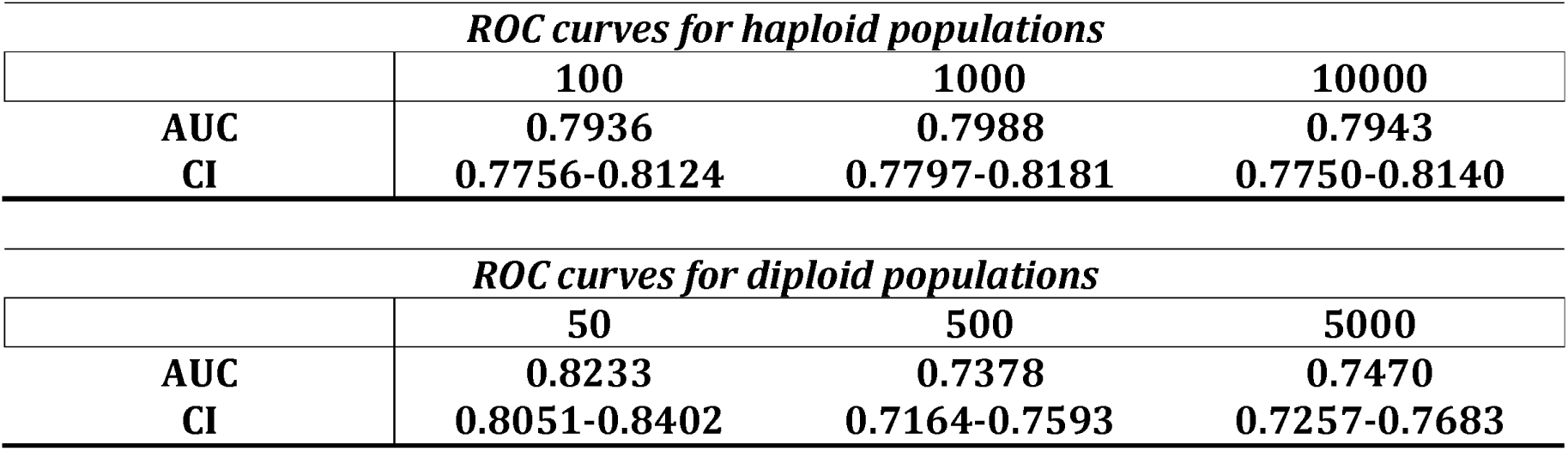
AUC values and confidence intervals for ROC curves.

**Table 2.** R^2^ values of the parameter estimation with the ABC model choice for haploid populations.

| *Case 1: 1000 constant selection pseudo-observables* |  |  |  |
| --- | --- | --- | --- |
| **M<sub>0</sub> (True)** | **100** | **1000** | **10000** |
| *s* | 0.558 | 0.843 | 0.914 |
| **M<sub>1</sub> (False)** | **100** | **1000** | **10000** |
| *s<sub>1</sub>* | 0.366 | 0.769 | 0.834 |
| *s<sub>2</sub>* | 0.608 | 0.752 | 0.685 |
| *CP* | -16 | -5.97 | -2.27 |

| *Case 2: 1000 changing selection pseudo-observables* |  |  |  |
| --- | --- | --- | --- |
| **M<sub>0</sub> (False)** | **100** | **1000** | **10000** |
| *s* | -0.278 | -0.00232 | -0.00205 |
| **M<sub>1</sub> (True)** | **100** | **1000** | **10000** |
| *s<sub>1</sub>* | -0.323 | 0.363 | 0.415 |
| *s<sub>2</sub>* | 0.704 | 0.768 | 0.777 |
| *CP* | 0.0946 | 0.413 | 0.303 |

**Table 3.** R^2^ values of the parameter estimation with the ABC model choice for diploid populations.

| *Case 1: 1000 constant selection pseudo-observables* |  |  |  |
| --- | --- | --- | --- |
| **M<sub>0</sub> (True)** | **50** | **500** | **5000** |
| *s* | -0.13 | 0.55 | 0.796 |
| *h* | -0.789 | 0.653 | 0.88 |
| **M<sub>1</sub> (False)** | **50** | **500** | **5000** |
| *s<sub>1</sub>* | -0.194 | 0.363 | 0.746 |
| *s<sub>2</sub>* | 0.491 | 0.57 | 0.609 |
| *h* | 0.195 | 0.764 | 0.857 |
| *CP* | -10.8 | -1.65 | -1.37 |

| *Case 2: 1000 changing selection pseudo-observables* |  |  |  |
| --- | --- | --- | --- |
| **M<sub>0</sub> (False)** | **50** | **500** | **5000** |
| *s* | -0.441 | 0.0955 | 0.118 |
| *h* | -0.119 | 0.525 | 0.573 |
| **M<sub>1</sub> (True)** | **50** | **500** | **5000** |
| *s<sub>1</sub>* | -0.713 | 0.181 | 0.277 |
| *s<sub>2</sub>* | 0.467 | 0.536 | 0.463 |
| *h* | 0.219 | 0.611 | 0.678 |
| *CP* | -0.145 | -0.258 | -0.125 |

Tables 2 and 3 summarize the performance of the joint parameter estimation for all cases as the *R^2^* values for the haploid and diploid populations, respectively. The first case is when the pseudo-observables are of constant selection intensity, in which case the true model (M_0_) performs only slightly better than the false model (M_1_) for estimating *s.* This discrepancy in parameter estimation of M_0_ is mainly owing to uninformative pseudo-observable trajectories with constant selection (which have been lost or fixed) being associated with the true model (M_0_) of constant selection; this is due to the constraint for the allele to be segregating at the change point in the (false) model M_1_ of changing selection. In the cross-validation of the constant selection case, the parameters estimated form horizontal lines at negative estimated values for those trajectories that are lost, and cluster at the top right corner for those trajectories that are fixed (Figure 3-4, Figure S4-S7).

For the second case in which the pseudo-observables have changing selection coefficients, parameter estimation from the true model (M_1_) performs better than that from the false model (M_0_) for all parameters – particularly when population sizes are large. As expected, there is a trend of better parameter estimation as population size increases. Additionally, it has been shown that the value of the Bayes factor *B_1,0_* is a good indicator of the parameter estimation performance (results not shown).

## Data Application

### Historical dataset of *Panaxia dominula*

A long-running dataset based on the *medionigra* morph responsible for darker wing color in wild populations of *Panaxia dominula* (Figure S8) began in 1939 with collections by Fisher (Fisher and Ford 1947) and continued through 1999 (Cook and Jones 1996; Jones 2000). Despite this phenotypic time-serial data having been analyzed previously from various angles (O’Hara 2005; Mathieson and McVean 2013; Foll et al. 2014b), it is still relevant to consider a model of changing selection in time, as Wright (1948) originally suggested.

The recent reconsiderations of the dataset tend to favor a lethal-recessive model with an effective population of 2*N_e_=* 1000 (Mathieson and McVean 2013; Foll et al. 2014b) – however, the biological question of how the *medionigra* morph could have reached the initial frequency of 11% in the dataset remains unanswered with this conclusion of constant strong negative selection. Wright asserted that the trajectory of the *medionigra* morph during this period could be explained by fluctuating selection with "no net selective advantage or disadvantage”. Although this alternative hypothesis has been considered as a random fluctuation of selection by estimating selection coefficients between every sampling time point (see O’Hara 2005), the quantitative plausibility of a directional change-in-*s* model over a single-*s* model lacks thorough investigation. Thus, we re-analyze this dataset using the CP-WFABC method in order to investigate the possibility of changing selection in the *medionigra* morph during the 60-year data collection.

Using the ABC model choice introduced here as a test for a change in selection strength, and to estimate the parameters of interest for the chosen model, we assume the *medionigra* allele is a single co-dominant locus responsible for the homozygous and heterozygous expressions of the phenotypic forms *bimacula* and *medionigra,* respectively (Cook and Jones 1996). The model M_0_ assumes a single selection coefficient, thus the only parameter to estimate is *s.* The M_1_ model assumes a change in selection strength, thus the parameters of interest are *s1, s2* and *CP.* Both M_0_ and M_1_ take the prior range of [-1,1] for the selection coefficients and the prior range of [2, 59] for the change point in the M_1_ model. For the M_1_ model, these uninformative priors are updated with the constraint that the allele must be segregating at the time of change point. Here, we create 10,000,000 simulated datasets for each M_0_ and M_1_, and apply the rejection algorithm of the CP-WFABC method to retain the best 1000 simulations compared with the observed trajectory. The effective population size is assumed to be 2*N_e_=* 1000 as in previous studies (Wright 1948; Cook and Jones 1996; O’Hara 2005), with an initial allele frequency of 11% and a minimum frequency ascertainment of 2%.

The Bayes factor for M_1_ over M_0_ is calculated as 0.952, indicating that the single coefficient M_0_ model cannot be rejected in favor of the changing selection M_1_ model (Table 4). From the parameter estimation of the model M_0_ (Figure S9), the mode of the posterior distribution for *s* is given as -0.15 as asserted by Fisher and Ford (1947). When the ABC model choice was repeated with a smaller population size of 2*N_e_=* 100 as suggested by Wright (1948) and O’Hara (2005), the Bayes factor increases to 1.87 (i.e., the changing selection model is twice as likely as the constant selection) – however, this value is not large enough to be significant for a diploid population of *N_e_ =* 50 (Table 4).

**Table 4.** The Bayes factor thresholds for the significance level α of 1% computed using the distribution of Bayes factors under the null model M_0_.

| *Diploid populations* |  |  |  |
| --- | --- | --- | --- |
| $\alpha=1\%$ | 50 | 500 | 5000 |
| BF | 4.7 | 1.6 | 1.3 |
| *Haploid populations* |  |  |  |
| $\alpha=1\%$ | 100 | 1000 | 10000 |
| BF | 3.7 | 3.2 | 1.7 |

### Experimental evolution of Influenza virus with drug treatment

The evolution of pathogens within a host is one of the most important cases in which the possibility of fluctuating selection must be considered – as they may experience drastically changing selective pressures due to host immune response, specific drug treatments, and/or pathogenic cooperation or competition (Tanaka and Valckenborgh 2011; Hall et al. 2011). Thus, how these pathogens adapt to these rapid external and internal changes is of major concern to the biomedical community.

The time-serial experimental dataset of influenza A conducted by Renzette et al. (2014) and Foll et al. (2014a) is an interesting case study on the impact of drug treatment on influenza virus evolution. The dataset consists of 13 sampling points from which population-level whole-genome data were collected. Drug treatment with a commonly used neuraminidase inhibitor (oseltamivir) began after the collection of the third sample and continued, at increasing concentrations, until the final passage. Using WFABC, the genome-wide effective population size across the sampling time points was estimated (*N_e_* = 176) and the SNPs under selection were identified.

Here, we apply the CP-WFABC method on two cases of interest from this study to consider a possible change in selection strength under drug treatment: the first case includes trajectories identified as being driven by positive selection, while the second includes outlier trajectories (i.e., trajectories not fitting a single *s* Wright-Fisher model). For all cases, we test the model M_0_ (i.e., a single selection coefficient) and M_1_ (i.e., a changing selection coefficient), with parameters of interest (*s*) and (*s1, s2, CP*), respectively. The number of generations per passage is assumed to be 13, and the minimum frequency of 2% is set as an ascertainment for observing the minor allele in the data. *De novo* mutations are assumed to occur at the first sampling time point for the SNPs whose allele frequency reached more than 2% before the drug administration (except for trajectories whose initial frequency is above 2%, which are assumed to be standing variation), and at the fourth sampling point for those whose frequency did not. This assumption is based on the high mutation rate, large population bottlenecks associated with passaging, and large census population size between passages. 10,000,000 datasets were simulated for each M_0_ and M_1_; the best 1000 trajectories from the lot were retained using the rejection algorithm described in the Methods. The uniform prior ranges for the selection coefficients were set as [-1,1], for the change point as [2,157] or [2,105] depending on the appearance of the mutation, and with the constraint of segregating alleles at the change point for M_1_.

The results for the Bayes factors and the parameter estimates are summarized in Table 5 for all trajectories of interest. The Bayes factors of most trajectories show strong support for the changing selection model: the stronger the selection strength change, the larger the Bayes factor. Using the Bayes factors from the simulation studies as guidance (Table 4), the significance threshold to reject M_0_ is computed as 3.7 for a small haploid population. As expected, the trajectories identified as outliers of the single *s* Wright-Fisher model (NP 159, PB1 33) all reject the constant selection model M_0_ with a large Bayes factor. We also note that the Bayes factor for the drug-resistant mutation H275Y (NA 823) does not support the changing selection model strongly, confirming that the experimental evolution procedure kept the selective pressure of the drug constant by adjusting the drug concentration to reduce viral plaque numbers to 50% at each passage. The change points are estimated to be mostly between the seventh and eighth passages, a notable result since three of these trajectories (HA 48, HA 1395, NA 582) are increasing rapidly after the drug-resistant mutation H275Y appears, whereas one trajectory (NP 159) from a different segment decreases rapidly. This result may indicate that positive selective for the three SNPs (including HA 1395; a known compensatory mutation encoded also as D112N) increased along with the drug-resistant mutation H275Y, potentially due to epistatic interactions, whereas another SNP decreased at that time, potentially owing to clonal interference. The single selection estimates from WFABC (Foll et al. 2014b) are similar to the M_0_ estimates of the constant selection coefficient only when the Bayes factor does not reject M_0_ – strong evidence that an alternative model of changing selection must be considered for some trajectories in order to correctly estimate selection coefficients.

**Table 5.**
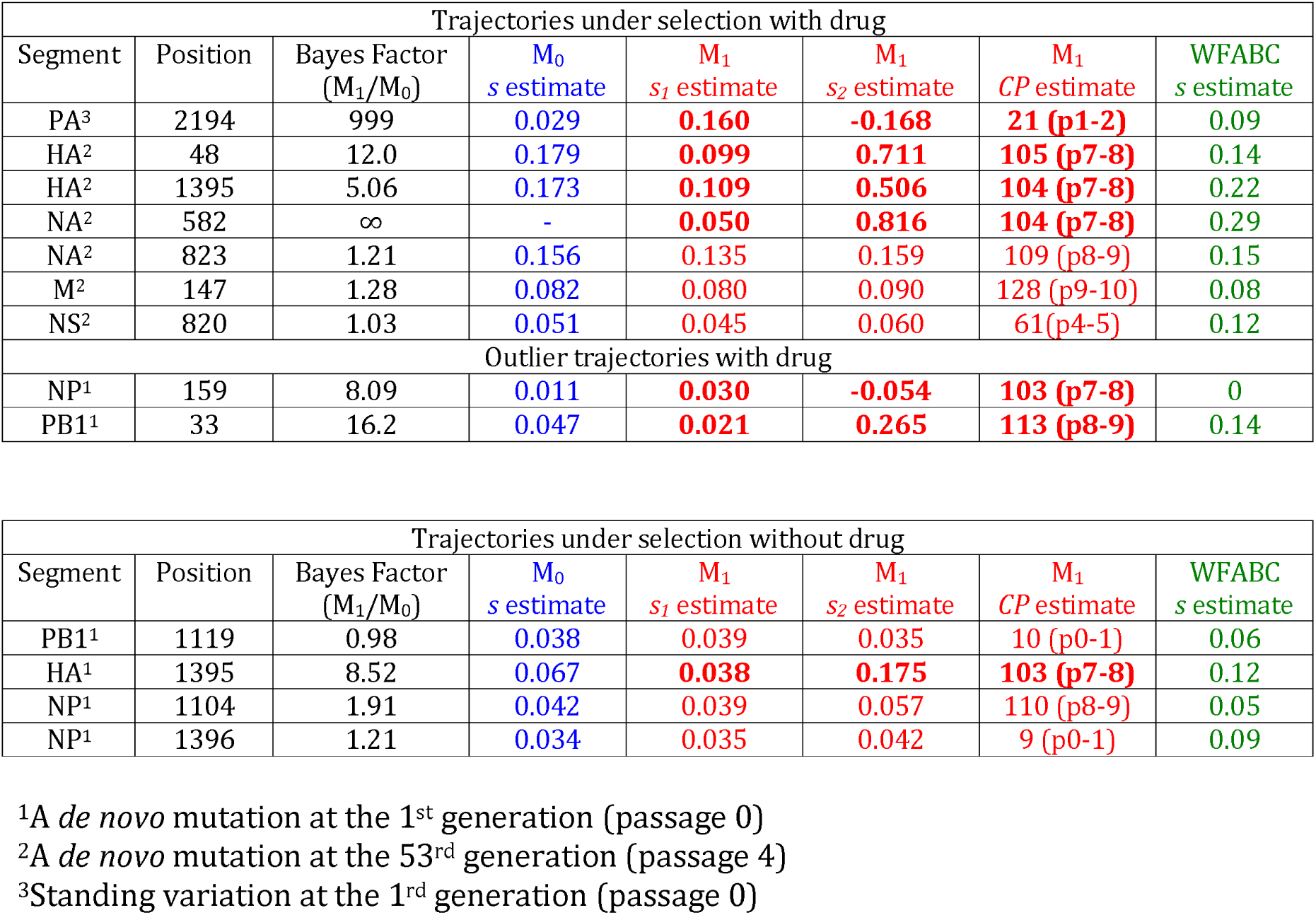
Bayes factors and parameters estimated for the influenza trajectories in the presence and absence of drug. The estimates whose Bayes factors show strong support for M_1_ are in bold.

We also applied CP-WFABC to the control case of SNPs increasing in frequency without drug as a comparison to the case with drug. The effective population sizes of the viral populations were averaged to be 226 in the absence of drug from the previous study (Foll et al. 2014a). For the control case, *de novo* mutations are assumed to occur at the first sampling time point for all SNPs, but the other inputs and the ABC model choice were kept the same as in the drug case. The Bayes factor results summarized in Table 5 demonstrate that three out of the four SNP trajectories under selection in the control experiment cannot reject M_0_ (i.e., constant selection). Interestingly, the only SNP trajectory to support M_1_ (i.e., changing selection) is HA 1395 – a known compensatory mutation that also appeared under drug treatment. The parameters estimated from the model chosen indicate there was a change in selective pressure from a slightly positive value to a strongly positive value between the seventh and eighth passage as shown in Figure 5b.

**Figure 5.**
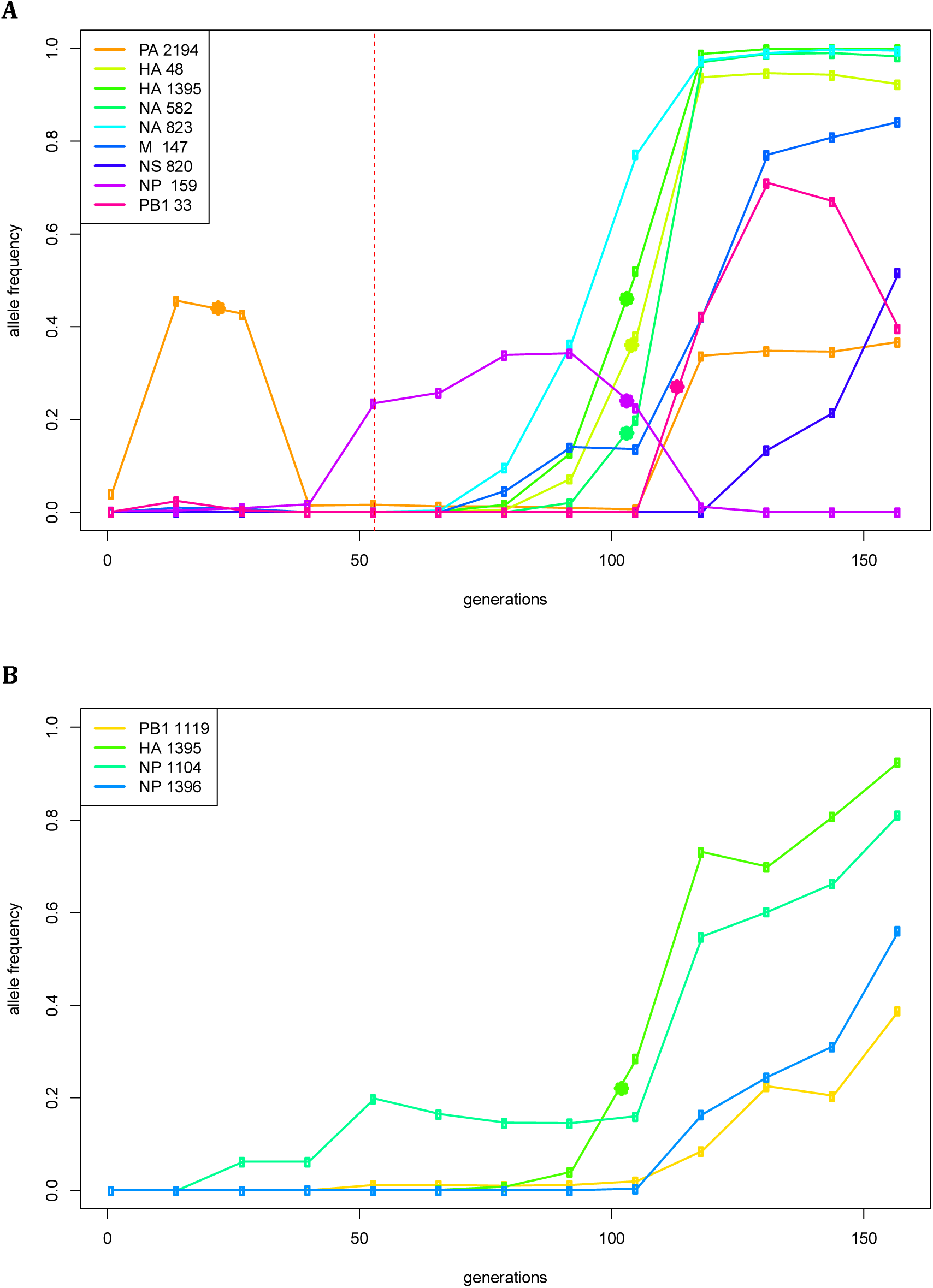
Change points indicated with solid stars for the trajectories of interest: (A) Increasing SNP trajectories in the presence of drug. The red vertical line indicates the sampling time of drug administration. (B) Increasing SNP trajectories in the absence of drugs.

## Discussion

These simulations demonstrate that the novel CP-WFABC approach presented here is able to detect changing selection trajectories via ABC model choice, and also to estimate a wide range of parameters of interest. Performance was analyzed separately for three categories of allele trajectories according to the nature of the change in selection strength: (1) a change from positive *s_1_* to positive *s_2_,* (2) a change from positive *s_1_* to negative *s_2_,* and (3) all other changes. The datasets for each possible combination were generated using the Wright-Fisher model with a change in selection strength, using the most general prior ranges for all parameters *s_1_, s_2_, CP,* and *h* for diploids, with the only constraint being segregation of the allele at the change point. For both the detection and parameter estimation, CP-WFABC performs the best when the change is large, particularly for the second category of change (positive *s_1_* to negative *s_2_*), as shown in the ROC curves (Figure 2, S3) and the cross-validation graphs (Figure 3-4, S4-S7). For the first category (positive *s_1_* to positive *s_2_*) and the third category (any other changes), the change point is difficult to estimate, particularly for diploids where the additional parameter *h* is also estimated. The ABC model choice of CP-WFABC has the best sensitivity for full specificity, for larger population sizes (*N_e_*>500 for diploids), and haploid populations.

The parameter estimates of *s_1_* and *s_2_* perform best when the values are moderate. For *s_1_,* the optimal parameter range for estimation is [0,0.5], where a *de novo* mutation that survives negative selection and segregates until the change point is indistinguishable from other drifting mutations with similar trajectories with uninformative low allele frequency. These trajectories naturally arise more frequently when population size is small and in diploids where the dominance effect plays a role, as shown in the third category of change (Figure 3-4, S4-S7; magenta points). When an initial frequency of 10% is used instead of a *de novo* mutation, the advantage of having a more informative trajectory at the beginning is counteracted by the effect of more cases under negative selection or genetic drift segregating until the change point. Thus, the performance of CP-WFABC for standing variation is similar to that of *de novo* mutation (results not shown). For *s_2_,* the optimal parameter range for estimation is [-0.5,0.5], as trajectories with extreme values are less informative since they are lost or fixed directly after the change point (explaining the clustering of the change points at earlier times). For diploid populations, estimates of *h* are accurate to the level of determining dominance from co-dominance or recessiveness, particularly for large population sizes (Figure S7), given the difficulty of joint estimation with the three other parameters. Indeed, estimation of this additional parameter comes at the cost of worse performance for the other parameters, as can be seen in the ROC curves and cross-validation graphs: the detection and parameter estimation of changing selection cases is always better for haploids. Thus, in diploid cases, we recommend fixing the dominance parameter if known, in order to improve the performance of CP-WFABC.

Although CP-WFABC is intended to detect and evaluate changing selection intensities, the simulation studies show that the method also performs well in estimating parameters for cases of constant selection – as has been demonstrated by Foll et al. (2014a). For haploids with large population sizes, in particular, the estimated values of the single parameter *s* correlate almost perfectly with the true values (Figure S5). However, when the population size is small for both diploids and haploids, some trajectories that are lost by negative selection or genetic drift are difficult to estimate, as shown in Figure 3 and Figure S6 as horizontal lines along some negative estimated values. This limitation of constant selection coefficients, however, is due to the simulation conditions of the *de novo* mutation at the first generation and the minimal ascertainment scheme (minimum frequency of 2% at one of the sampling time points). For real datasets, the conditions are likely to be less stringent, since such uninformative trajectories will not be considered for parameter estimation.

Finally, we utilized this approach to make inference in two very different time-sampled datasets: *Panaxia dominula* (diploid) and Influenza A (haploid). The time-serial *medionigra* trajectory of *P. dominula* was re-analyzed to test for a change in selection strength and/or direction during 60 years of data collection. By assuming *h* as codominant, the results for *N_e_ =* 500 indicate that the model M_0_ of constant selection cannot be rejected according to the Bayes factor from the ABC model choice algorithm. The selection coefficient from this model is estimated as -0.15, corresponding with that calculated by Fisher and Ford (1947). However, when the population size is assumed to be smaller (*N_e_ =* 50), the Bayes factor result supports M_1_ (changing selection) twice as strongly as M_0_ (constant selection), but not strong enough to reject M_0_ according to the significance level test computed with the distribution of Bayes factors under the null model M_0_. This dataset of the *medionigra* morph thus demonstrates the difficulty of detecting and evaluating a change in selection when the population size and the number of generations are small.

Next, CP-WFABC was applied to SNP trajectories of interest from an experimental dataset of influenza A virus. The ABC model choice test was conducted on the trajectories identified as 1) being positively selected and 2) as outliers from the single-*s* WFABC method. For the SNPs in the presence of drug, the Bayes factor for six out of nine trajectories favored the changing selection model M_1_. The change points for four out of these six trajectories occurred between passages 7 and 8 – the interval during which three trajectories from the segments HA and NA increased rapidly while one trajectory from the segment NP decreased rapidly along with the known drug-resistant mutation NA 823 (H275Y). These results appear to support the presence of epistasis and clonal interference, where the selection strength of the other SNPs is influenced by the appearance of a drug-resistant mutation under drug pressure. In fact, a known compensatory mutation (HA 1395) was among the three trajectories increasing rapidly, reinforcing the use of the method to evaluate biological hypotheses. Moreover, the estimated values of selection coefficients differed greatly between the constant-selection method (WFABC) and CP-WFABC. In particular, the estimates of *s* for outlier trajectories from the Wright-Fisher model were inferred by WFABC as being near zero (neutral), though the more robust CP-WFABC estimates here indicate fluctuation of *s* from negative to positive values. Thus, these fluctuations cannot be explained by genetic drift alone, as previously speculated. We have therefore identified some cases where an alternative model of changing selection is essential for correctly estimating selection parameters and identifying change points. For the SNPs in the absence of drug treatment, the Bayes factor for three out of four trajectories could not reject the constant selection model M_0_. This result indicates that in the absence of drug, the selective pressures on the population are largely constant as expected. The SNP trajectory identified as supporting M_1_ in the control case is a known compensatory mutation (D112N) for infectivity (Thoennes et al. 2008) that also appeared in the presence of drug, further confirming the increased infectivity might contribute to the tissue culture adaptation.

The simulation studies of changing selection reveal some important points to consider from the standpoint of experimental design. Firstly, at least two sampling time points are needed to estimate selection strength in time-serial methods. For CP-WFABC, the parameter estimation of a single selection coefficient between two sampling time points performs reasonably well for haploid population sizes above *N_e_*=1000. However, it is advisable to have three sampling time points to maximize the performance of parameter estimation, particularly for diploids and in smaller population sizes (*N_e_*<1000). Thus, in order for a change to be detectable, it is required to have at least four sampling time points where the change must occur between the second and the third sampling time points – an important factor to consider for the design of change-point experiments, such as drug administration or environmental change. The simulation studies of CP-WFABC confirm that the estimation of *CP* performs best at the intermediate range of time-sampled data, as any change happening before the second and after the second-to-last sampling time point is impossible to detect (Figure 3-4, S4-S7). Finally, it remains a future challenge to expand this method to consider more than one change point in selection strength, as some of the trajectories in the influenza A application (such as PA 2194 and PB1 33) suggest the presence of several change points along the trajectory.

## Acknowledgements

We thank Kristen Irwin and Sebastian Matuszewski for helpful comments. The computations were performed at Vital-IT (http://www.vital-it.ch) in the Swiss Institute of Bioinformatics (SIB). This work was funded by grants from the Swiss National Science Foundation and a European Research Council (ERC) starting grant to JDJ.

## Data Accessibility

The R package of CP-WFABC is available on jensenlab.epfl.ch.

